# Teddy: neural inference of epidemiological parameters from viral sequences

**DOI:** 10.64898/2026.01.05.697728

**Authors:** Vincent Garot, Luc Blassel, Luca Nesterenko, Anna Zhukova, Samuel Alizon, Laurent Jacob

## Abstract

Estimating how fast infections spread or how long they last is essential to control outbreaks. Phylodynamics methods enable the inference of key epidemiological parameters from viral genomic data but remain limited in terms of biological realism and speed because they need to derive and compute likelihoods. We address this issue using simulation-based inference and introduce a deep learning–based framework directly trained on alignments of viral genetic sequences. Our neural posterior estimation matches the accuracy of leading Bayesian likelihood-based methods while running a thousand times faster and avoiding a phylogeny reconstruction step. This performance can be harnessed to analyze large datasets and opens new perspectives to tackle biologically realistic models in terms of pathogen life histories or genomic evolution.

---

The timely and accurate estimation of outbreak parameters is of key importance for public health and a cornerstone of quantitative epidemiology [1]. This classically relies on inci-dence data (*i.e.*, the number of new cases in the population per unit of time), as well as on patient monitoring or contact-tracing data, which both require costly field studies.

Progress in molecular and computational biology has allowed the use of microbial genetic sequences instead of incidence data to infer these parameters through the field of phylodynamics, which hypothesizes that such inference is possible by combining population dynamics models and phylogenetics [16, 43, 11]. The underlying assumption is that phylogenetic trees, *i.e.*, virus genealogies, provide information about past transmission events. Phylodynamics involves two inference tasks: estimating virus phylogenetic trees from sequences to approximate transmission trees and estimating epidemiological parameters from these trees. In practice, using only a hundred dated virus sequences, phylodynamic methods can accurately estimate epidemiological parameters such as the basic reproduction number, classically denoted *R*_0_, which corresponds to the average number of secondary infections an infected individual generates in a fully susceptible population, or the mean duration of the infectious period, here denoted *δ* [34, 42].

Current approaches suffer from two important limitations. First, they typically assume independence between genome evolution and virus transmission, thereby ignoring important biological processes such as recombination, selection, or within-host virus diversity. Second, they typically require computations of likelihood functions that correspond to the evolutionary and epidemiological models, which has been a strong restriction on the architecture, and therefore on the realism, of the models. For example, frequent phenomena such as co-evolution between sites in sequence evolution or multiple infections in population dynamics already lead to intractable likelihoods. Furthermore, even simplified models such as the susceptible infected susceptible (SIS) are challenging to derive and lead to costly computation [23], thereby generating a trade-off between realism and affordability, which eventually limits our ability to accurately estimate epidemiological parameters.

Simulation-based inference (SBI, [7]) has emerged as a powerful paradigm to handle probabilistic models with intractable likelihoods. Its key idea is to use simulations to generate observations rather than deriving and evaluating a likelihood function. In many cases, *e.g.*, here for population dynamics and sequence evolution models, simulation can be a rapid operation when a formal evaluation is intractable. One of these approaches is Neural Posterior Estimation (NPE, [25]), which consists in training a neural network over samples from the probabilistic model. The resulting networks can formally be shown to approximate the posterior distribution under this model [30]. When it comes to infer-ring epidemiological parameters from phylogenies, NPE can achieve comparable results as state-of-the-art likelihood-based Bayesian inference methods, without incurring the compu-tational cost [44]. Similarly, for the upstream task of inferring phylogenies from sequences, recent work shows that NPE can match or even outperform the accuracy of likelihood-based approaches for a fraction of the computational time, with the possibility to explore probabilistic models of sequence evolution that are intractable using likelihood-based methods [28, 3].

Building on these earlier studies, we introduce Teddy, an NPE approach for inferring epidemiological parameters from a set of dated viral sequences in a single step, *i.e.*, without inferring a phylogeny. We hypothesized that since current neural architectures can accurately perform phylogenetic reconstruction, they should also capture all necessary information about the tree from the sequences. Assuming tractable models for epidemi-ological dynamics and for sequence evolution, we demonstrate that Teddy is at least as accurate as the state-of-the-art likelihood-based inference software package BEAST 2 [4] to infer *R*_0_ and *δ*. Strikingly, the estimates made by the two approaches are highly correlated.

We also verify that the two yield consistent estimates for an alignment of hepatitis C virus (HCV) genomes from a past outbreak. Finally, since Teddy runs three orders of magnitude faster than BEAST 2, as it never needs to evaluate a likelihood, we show that this speed can be leveraged to analyze large datasets via subsampling procedures. Overall, since it does not require a decoupling between population dynamics and sequence evolution and relies on simulations, Teddy paves the way for fast inference assuming biologically realistic models of infectious disease spread, microbial genome evolution, and their interaction.

## Results

### Likelihood-free and phylogeny-free inference

Our Transformer for EpiDemiological DYnamics (Teddy) is summarised in Figure 1. It is a learnable function that estimates quantiles of the posterior distribution of virus population dynamics parameters, here the basic reproduction number *R*_0_ and the infection duration *δ*, from input data, here a dated multiple sequence alignment (MSA) and the value of its sampling proportion *s*. We briefly explain its structure and development, additional details are available in the Methods section.

**Figure 1:**
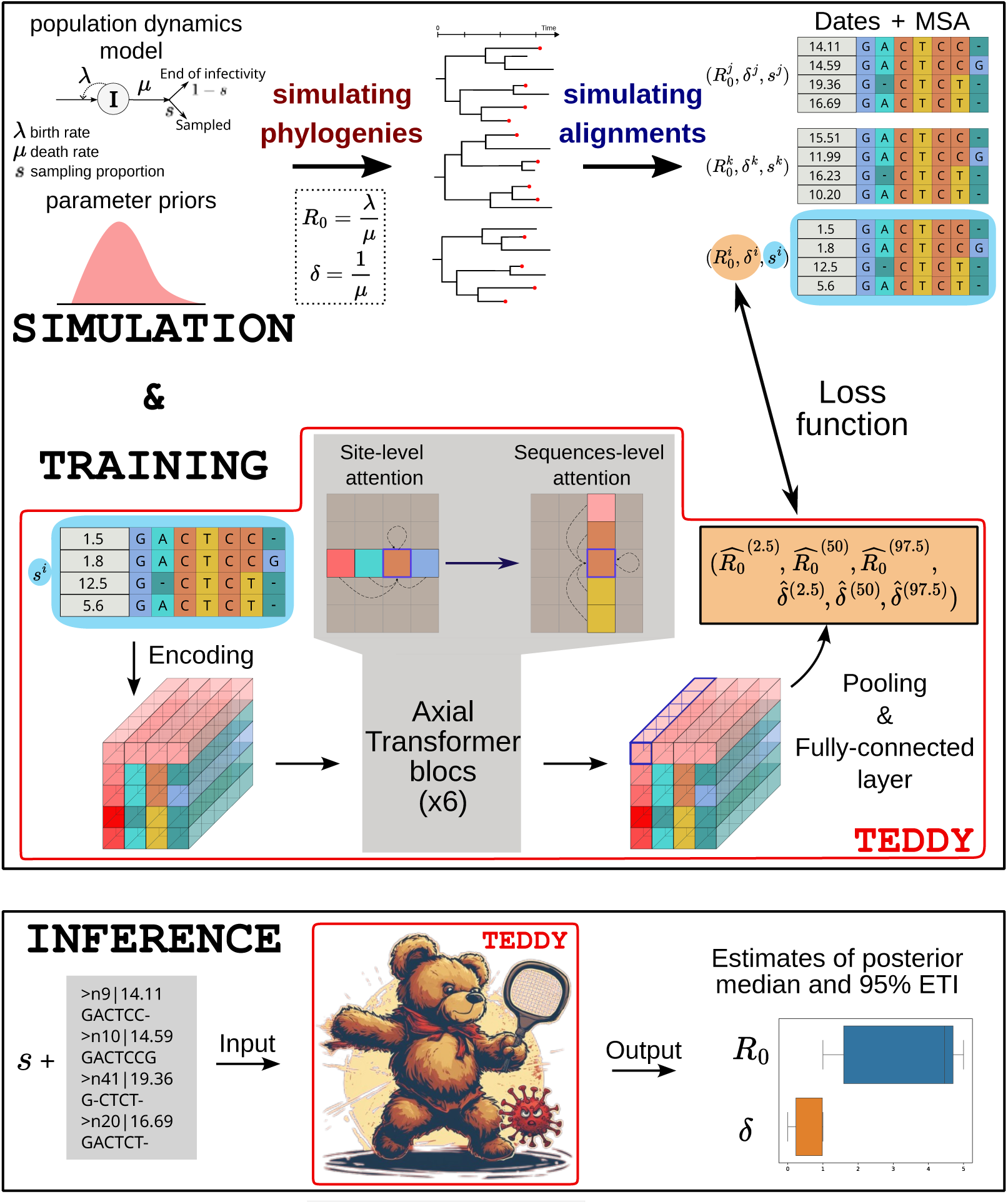
Schematic of the Teddy workflow. Teddy learns an approximation of the posterior median and Equal-Tailed credible Interval (ETI) for epidemiological parameters from samples of a probabilistic model. We generate pairs of epidemiological parameters and dated multiple sequence alignments (MSA) by sampling the parameters from a prior distribution, simulating phylogenies from a population dynamics model using these param-eters, and simulating alignments from a probabilistic model of sequence evolution along these phylogenies (top). The Teddy neural network transforms the dated alignment into quantile estimates through axial transformer_7_blocks. We learn the weights of the networks using the simulated pairs (epidemiological parameters, dated MSAs). The loss function used to compare the network outputs and simulated parameters ensures that they approx-imate the posterior quantiles (middle). For the inference, we apply the trained Teddy architecture to a dated MSA and use the outputs to estimate the posterior distribution median and ETI (bottom).

To train our algorithm, we simulate input data from a conditional probabilistic model, the likelihood of which is denoted *p*(dated MSA|*R*_0_*, δ, s*), and sample the values of the epidemiological parameters to estimate (*R*_0_*, δ*) from a prior distribution. The input sampling proportions *s* are chosen to cover a large epidemiologically plausible interval.

Teddy starts from a raw representation of an MSA where the embedding of each site within each sequence only encodes its own nucleotide. The sampling dates and the sampling proportion *s* are set and encoded as side information. We then apply a series of transformations (namely axial attention blocks [31]) that allow information to flow between sites for each sequence and between sequences for each site. Following these transformations, every entry in the array captures information about the whole alignment. This enriched representation is used to generate predictions for the epidemiological parameters, denoted (*R*^^^_0_*, δ*^^^).

Finally, we optimize the learnable weights of Teddy to make its output (*R*^^^_0_*, δ*^^^) as close as possible to the parameter values (*R*_0_*, δ*) used for simulation of each dated MSA in the training set. For this, we use an appropriate loss function such that the resulting trained network approximates quantiles of our posterior distribution of interest, namely *p*(*R*_0_*, δ*|dated MSA*, s*). We expect a successfully trained Teddy to estimate quantiles of the posterior as accurately as a Markov chain Monte Carlo (MCMC) algorithm, but without requiring the (computationally costly) evaluation of the likelihood function *p*(dated MSA|*R*_0_*, δ, s*). Note that although in Figure 1 a phylogeny is drawn from a population dynamics model before an alignment is drawn from a separate model along this phylogeny, this need not be the case. As long as the simulator is fast enough, sequence data can be directly generated from the population dynamics model, thereby allowing for processes such as recombination or multiple infections to take place.

### Fast and accurate inference

We assessed the performance of Teddy by comparing its performance to that of BEAST 2 for a Birth-Death-Sampling epidemiological model (BDS, [33]). We chose this model because it has a closed-form solution for its likelihood function and hence does not require numerical approximations during parameter estimation in BEAST 2 (see Supplementary Text S1 and Figure S6). For the nucleotide sequence evolution, we chose the Jukes-Cantor model (JC, [20]) for its simplicity.

We simulated 1, 500, 000 MSAs under the BDS-JC setting for training Teddy, and an additional 1, 000 MSAs to compare the methods. Using only 500, 000 led to similar results but proved more sensitive to the number of training steps (Supplementary Figure S4). The distributions of the parameters used to generate the training and test data are given in the Methods. We compared the Mean Absolute Error (MAE), the Mean Relative Error (MRE), the coverage (*i.e.*, the percentage of cases where the true parameter value lies in the estimated credible intervals), and the mean width of the credible intervals—see Methods for more details. The lower the MAE, the MRE, and the mean width, the better. Similarly, the closer the coverage is to 95 % the better.

Teddy showed better results than BEAST 2 in terms of MAE, coverage, and mean width of the Equal-Tailed credible Interval (ETI), while the MRE was lower for BEAST 2 (Table 1). The MAE and MRE results are consistent with the fact that Teddy was trained to minimize the former—whose true minimizer over the posterior distribution is the actual median. The slightly larger MRE indicates that Teddy tends to make more errors for small parameter values than BEAST 2, an issue that could be addressed by changing its objective. On the other hand, Teddy has better coverage than BEAST 2, even though it has narrower credible intervals. This implies that it is better calibrated than BEAST 2 for quantile estimations.

**Table 1:** Average performance of BEAST 2 and Teddy on simulated data. Mean Absolute Error (MAE) and Mean Relative Error (MRE) are computed over 1, 000 test MSAs. The coverage corresponds to the percentage of true parameters in the predicted 95% Equal-Tailed credible Interval (ETI). We also compute the Mean Width of the estimated ETI over the test set. For the MAE, MRE and the Mean Width ETI, we computed the standard error of the estimator of the mean (sem). For each summary statistic, the method with the best performance is shown in bold.

| | $R_0$ | | $\delta$ | |
| --- | --- | --- | --- | --- |
|  | BEAST 2 | Teddy<sub>0</sub> | BEAST 2 | Teddy<sub>0</sub> |
| MAE (sem) | 0.495 (0.014) | **0.488** (0.013) | 1.09 (0.03) | **1.07** (0.03) |
| MRE (sem) | **0.170** (0.005) | 0.174 (0.006) | **0.243** (0.008) | 0.245 (0.009) |
| % in ETI | 94.2 | **95.2** | 92.4 | **95.1** |
| Mean Width ETI (sem) | 2.10 (0.03) | **2.09** (0.03) | 4.62 (0.06) | **4.59** (0.05) |

The predictions from BEAST 2 and Teddy were highly correlated for the median and the quantiles (Figure 2). This is consistent with Teddy estimating the same Bayesian posterior distribution as BEAST 2—only without computing likelihoods.

**Figure 2:**
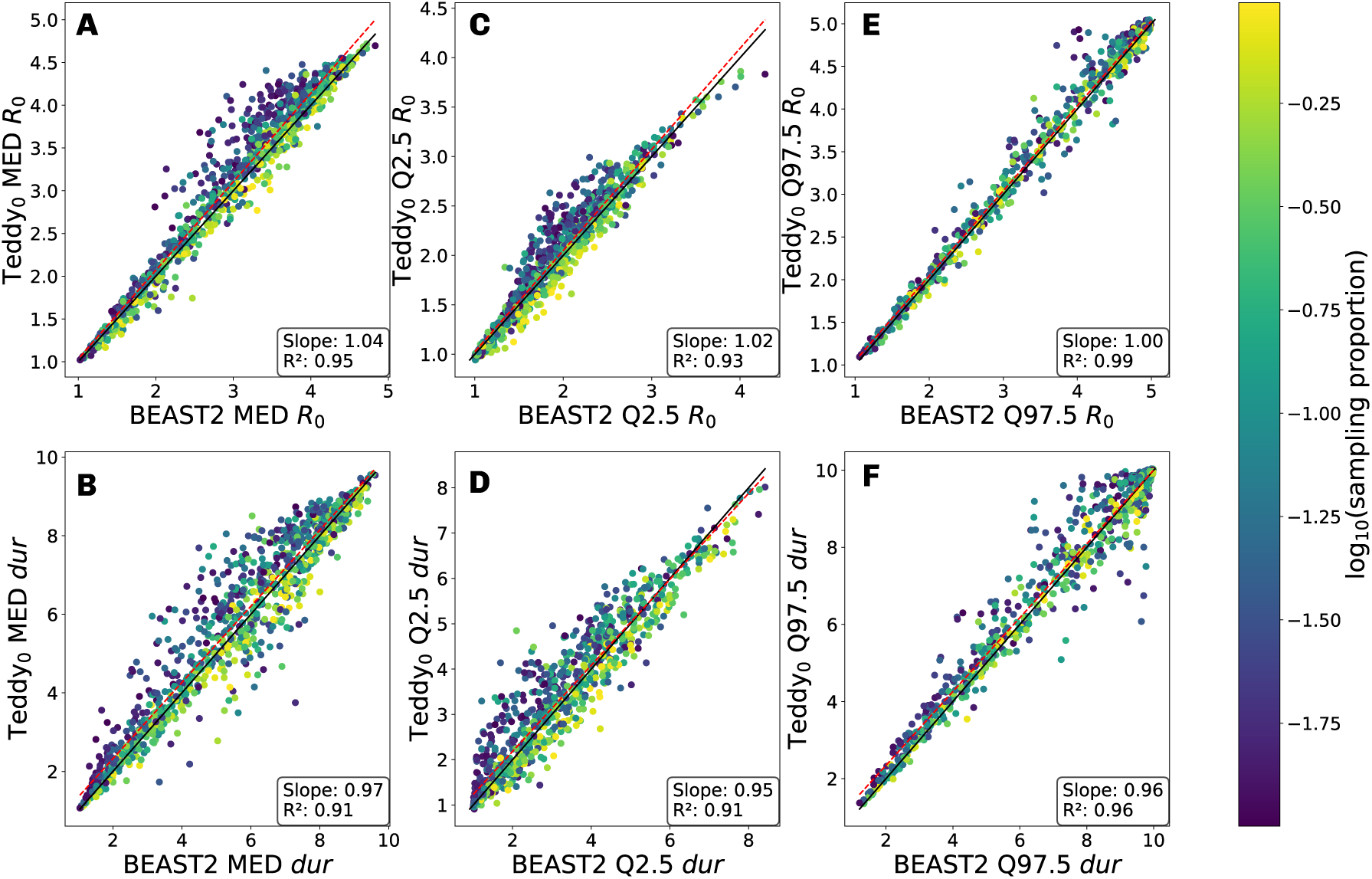
Estimations of Teddy_0_ versus BEAST 2 for the quantiles of the population dynamics model parameters. X-axes show the estimate from BEAST 2 and y-axes that from Teddy_0_. Panels A, C, and E show the estimates for *R*_0_ and panels B, D, and F for infection duration (*δ*). Panels A and B show the median, C and D the 2.5 % quantile, and E and F the 97.5 % quantile. Each point corresponds to the estimate on one of the 1, 000 test MSAs generated under the BDS-JC model described in the Methods. Colors indicate the value of the sampling proportion parameter *s*. The bottom-right boxes on each subplot and the dashed red lines show the estimates of a linear regression.

In our experiment, BEAST 2 provided more accurate inferences in runs with higher Effective Sample Sizes (ESS) [14] of the MCMC (Supplementary Figure S1). This hints that BEAST 2 does progressively converge towards the true posterior distribution and that longer MCMC chains could improve BEAST 2 accuracy, although the associated increase in computation time could become prohibitive. However, BEAST 2 convergence is usually assumed to be reached as soon as the ESS is greater than 200, and our results show that BEAST 2 does not outperform Teddy until ESS values become much higher.

In terms of computation speed, once trained, Teddy performed the inference for 100 test MSAs within about 2.5 minutes on a CPU—or 2.5 seconds on a single GPU. In comparison, it took BEAST 2 15 minutes on average to handle a single MSA on a CPU. Training Teddy took 19 hours on 4 H100 GPUs, but it is only necessary once for each probabilistic model and parameter range. Re-training or fine-tuning may be needed only if the epidemiological model, the evolutionary model, or the prior distribution are changed. In the following, we refer to this trained version of Teddy as Teddy_0_.

### Consistent performance on real outbreak data

We assessed the performance of Teddy on a real-life alignment of 35 sequences obtained from a Hepatitis C virus (HCV) outbreak in Bangkok, Thailand [18]. HCV has a high mutation rate and a very low recombination rate, making it well-suited to the phylodynamic framework and the site-independent sequence evolution model, respectively. Since we lack *R*_0_ and infection duration *δ* field estimates for this outbreak, our main objective was to verify that Teddy’s inference is consistent with that of BEAST 2 when analyzing virus genetic data.

For BEAST 2, following [8], we assumed a more realistic model of sequence evolution and a wider prior on the duration of infection than in the Teddy_0_ setting (see the Methods subsubsection on real outbreak data for more details). Before analyzing the real HCV alignment, we compared the performances of BEAST 2 and Teddy_0_ on 1, 000 alignments simulated under this new setting, using the same metrics as in Table 1. All these align-ments were simulated by starting from an HCV sequence from the outbreak. As expected, the inference provided by BEAST 2—under the same model used to simulate the 1, 000 alignments—was markedly different from the one obtained with Teddy_0_, which was trained under a different evolutionary model and a narrow epidemiological prior (see Supplemen-tary Figure S2). Supplementary Table 1 further shows that the inference provided by BEAST 2 is more accurate than the one obtained from the misspecified Teddy_0_.

Therefore, we re-trained Teddy on a new set of alignments generated under the same prior and model of sequence evolution as BEAST 2 and our simulated data—including the root sequence. As in Table 1, this new Teddy_HCV_ yielded more accurate estimates than BEAST 2 on the 1, 000 test alignments generated under the same model (right column of Supplementary Table 1). The Teddy_HCV_ and BEAST 2 estimates were very correlated (Supplementary Figure S3), which is expected since both methods perform Bayesian inference under the same probabilistic model. Fine-tuning, *i.e.*, starting this new training from the weights of the initial Teddy (Teddy_0_) instead of training from scratch allowed for faster convergence, implying that adapting Teddy to a new model can be less expensive than the initial training.

Finally, we compared the inferences of Teddy_HCV_ and BEAST 2 on the 35 HCV sequence alignment. Since the sampling proportion was not known, we show the results for two scenarios (*s* = 0.1 and *s* = 0.5, Figure 3). The two estimators yield similar results for *R*_0_ and the duration of infection *δ*, independently of the sampling proportion, which itself can affect the estimates of (*R*_0_*, δ*). The magnitude of the estimates, especially the duration of infectiousness of approximately 4 years, are comparable with earlier studies on different HCV outbreaks in Egypt and in France [33, 8].

**Figure 3:**
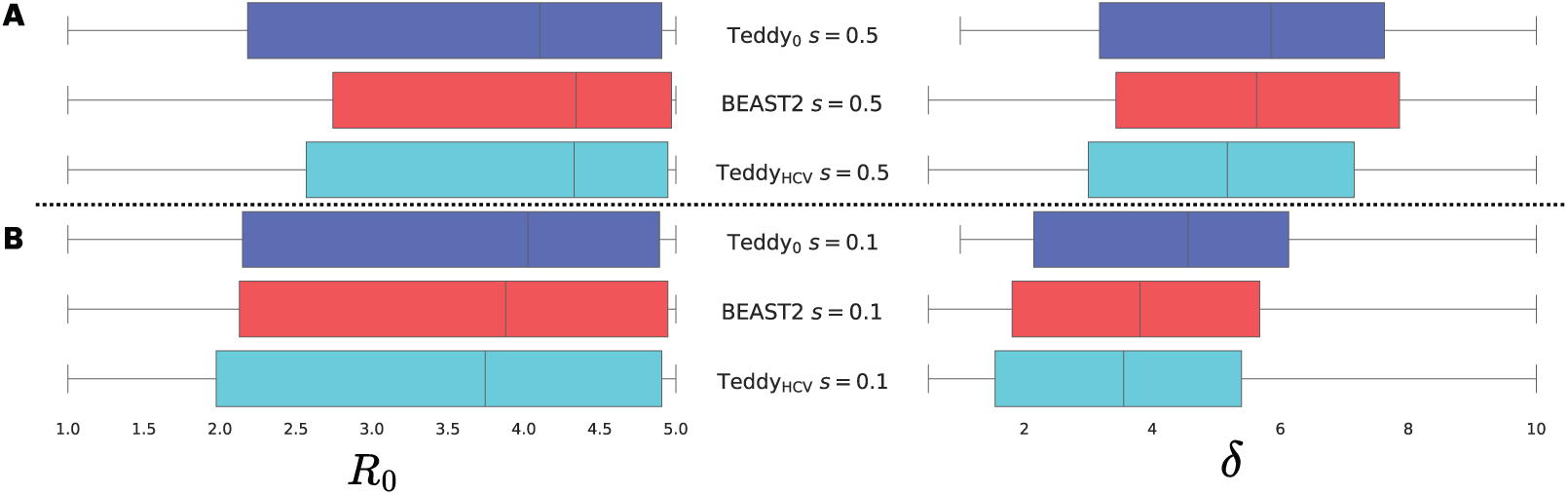
Infering epidemiological parameters on sequence data from a HCV outbreak in Bangkok (Thailand). Inferences are done using BEAST 2 (in red), Teddy_0_ (before fine-tuning, in blue) and Teddy_HCV_ (fine-tuned, in cyan). Panel A assumes a high sampling proportion (*s* = 0.5) and panel B an intermediate sampling proportion (*s* = 0.1). The priors in Teddy_HCV_ are identical to those of BEAST 2. The left column corresponds to *R*_0_, while the right – to infection duration *δ*. The boxplots cover the estimated ETIs (with the median indicated), while their whiskers show the prior intervals (which are shorter for Teddy_0_ in the case of *δ*). The HCV dataset contains 35 sequences and is further described in the Methods.

### Subsampling to accurately analyse large alignments

Teddy can currently be trained on alignments of up to 6 000 sequences with 250 variable sites, and its computation time and memory scale linearly with the product of the number of sequences and sites. Since the inference requires less memory than the training, a trained Teddy model can be applied to large alignments of at least 50 000 sequences with 250 variable sites. However, the estimation accuracy decreases as we move away from the sizes of its training examples, a phenomenon commonly known as length overfitting in the Transformers literature [2]. For example, when using Teddy_0_, which was trained on MSAs of 20 to 100 sequences, to analyze MSAs of more sequences, we obtain higher absolute errors than with MSA of 100 sequences (blue line in Figure 4).

**Figure 4:**
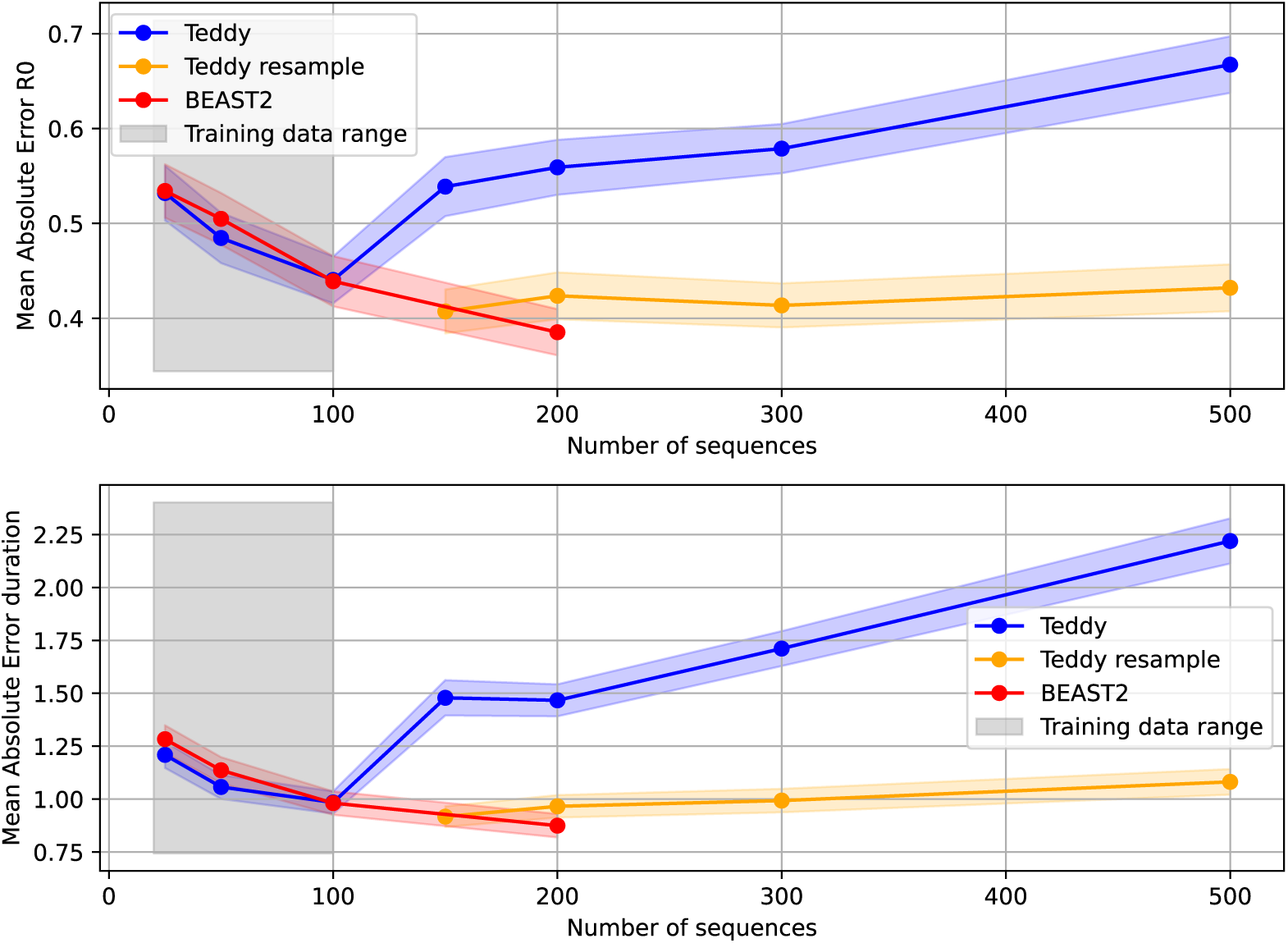
Mean Absolute Errors for estimates of *R*_0_ **(top) and duration (bottom) obtained from Teddy_0_ and its ensembling version on similated data.** Means are computed over 1, 000 MSAs with variable numbers of sequences (indicated on the x-axis), each generated under the model described in the Methods. The ensembling version, denoted ‘resample’ (in orange), draws 10 subsets of 100 sequences from the dataset, infers parameters with Teddy_0_ on each subset, and returns the median of the estimates. It performed better than trying to use all of the sequences (in blue) and is comparable to Beast 2 (in red). Shaded areas around curves correspond to 1.96 standard error intervals. The gray shaded area highlights the training data zone for Teddy_0_.

However, thanks to the speed of Teddy’s inference, we can readily apply it to multiple subsamples of the data of sizes consistent with the training dataset. This avoids retraining the algorithm, which is computationally demanding. In the case of our MSA of 200 sequences, running Teddy_0_ on ten subsamples of size 100 sequences and taking the median of the estimates yields absolute errors compatible with the ones reached by BEAST 2 on the 200 sequences (violin plot in Supplementary Figure S5). This subsampling can be per-formed on larger datasets and we show in Figure 4 that our ensembling strategy remains accurate with up to 500 sequences in the target dataset. However, the larger the dataset, the more difficult the comparison with BEAST 2 because of its computational cost. With regards to the alignment length, as described in the Methods, we already perform this sampling by default if the MSA has more than 250 variable sites. Future work will explore these strategies to improve the scaling of Teddy to larger alignments.

## Discussion

Inferring population dynamics parameters from sequence alignments is a challenging task. We introduced Teddy, a fast likelihood-free method to infer epidemiological parameters that uses virus genomic sequence data as input, without requiring any phylogenetic reconstruction. When assuming a tractable homogeneous Birth-Death-Sampling epidemiological model and a standard sequence evolution model, Teddy’s inference was at least as accurate as BEAST 2’s, a standard likelihood-based Bayesian inference tool, while running three orders of magnitude faster.

As for most existing methods, Teddy becomes limited for large datasets. However, its rapidity makes it possible to readily subsample the dataset in order to generate hundreds of smaller datasets of sizes compatible with that used to train the algorithm. We show that such an ensemble approach, which returns a distribution of the median value of the estimate of each of the small datasets, provides results comparable to likelihood-based methods. Such a scalable method can prove to be particularly useful given the increase in genetic data availability. Future work will explore the optimal subset size in terms of accuracy and computational efficiency.

Teddy and BEAST 2 provided similar inferences on a dataset from an HCV outbreak. For Teddy, this task required adapting a pre-trained model to the biology of this virus and its spread, which was more computationally frugal than training a new model from scratch. Note that retraining is only necessary when analyzing an outbreak caused by another virus (*e.g.*, SARS-CoV-2 instead of HCV) or in a population with characteristics that fall out of the priors (which is unlikely because these are large). For most MSAs originating from HCV outbreaks, Teddy_HCV_ can be used and provide results in a matter of seconds.

The current Teddy architecture correctly estimates quantiles of the posterior distributions, but a future goal will be to approximate the full posterior distribution, *e.g.*, by using normalizing flows [29]. Even though practitioners are mainly interested in credible intervals, working with joint distributions of parameters will provide access to their correlation structure and help handle non-identifiable probabilistic models.

One of Teddy’s current limitations is that it is unable to quantify epistemic uncertainty [21], and in particular to detect model misspecification or out-of-distribution (OOD) samples. It does not have measures of convergence (such as the ESS in BEAST 2), and data subspaces of very low prior probability are by definition undersampled, which implies that estimations with Teddy in these subspaces can be less accurate. One possible extension for Teddy to estimate these uncertainties would be to lean on OOD detection [19], a field that is largely developed for computer vision [45], or on other techniques such as dropout for Bayesian approximations [13].

Finally, Teddy’s full potential as a likelihood-free method will be unlocked when per-forming inference under probabilistic models with intractable likelihoods. For example, high-resolution spatial epidemic models can currently be used for simulations but not for inference (*e.g.*, [38]). Furthermore, more realistic sequence evolution models involving re-combination can make phylogenetic inference extremely challenging. Methods using phylogenetic networks have already been explored [27], but tend to be slow and remain limited. Another reason why the assumption that a phylogenetic tree can approximate a transmission tree may be unwise is that it does not take into account within-host virus diversity and neglects the cotransmission of different genotypes [17]. In all these situations, Teddy trained under the appropriate simulations could offer more realistic estimates than those currently possible under tractable models, and improve our ability to accurately inform public health policies.

## Methods

### Simulation-Based-Inference

In this work, we leverage recent advances in the simulation-based (or likelihood-free) in-ference paradigm [7], and, more specifically, on neural posterior estimation (NPE, [25]). NPE relies on a neural network Ψ(*x*) to parameterize an approximation *q*_Ψ(_*_x_*_)_(*θ*|*x*) of the posterior distribution of parameter *θ* given observed data *x*. The neural network is trained to maximize some function of *q*_Ψ(_*_x_*_)_(*θ*|*x*) over examples of (*x, θ*) generated by successively sampling *θ* under a prior *p*(*θ*) and *x* under a probabilistic model *p*(*x*|*θ*). Depending on the choice of the maximized function, the *q*_Ψ_ function based on the trained network can be shown to formally approximate the posterior *p*(*θ*|*x*) = *p*(*x*|*θ*)*p*(*θ*)*/p*(*x*) or some quantities depending on this distribution [30]. In this work, we minimize the Pinball loss (see below) and the resulting network approximates quantiles of the posterior. NPE has gained traction in evolutionary genomics because it only accesses *p*(*x*|*θ*) through sampling, thereby allowing inference under probabilistic models whose likelihood is expensive or impossible to evaluate. Another important feature of NPE is that the approximation is amortized: training the network can take time and resources, but, once trained, performing inference on a new data *x*_0_ is typically much faster than with likelihood-based approaches (assuming these can be available).

### Transmission and evolution models

To allow the comparison with likelihood-based inference approaches, we assume that the underlying epidemiological dynamics follows a simple Birth-Death-Sampling (BDS) model [33], which is often used to approximate the Susceptible-Infectious-Recovered (SIR) model [1] in infectious disease epidemiology. In this model, an infectious person generates a new in-fection at a rate *λ* and becomes non-infectious at a removal rate *µ*. For a proportion *s* of these ‘removed’ individuals, the microbe causing the infection is sampled and its genome becomes available for phylodynamic analysis. The basic reproduction number, which corre-sponds to the average number of secondary infections generated by an infected host during their whole infectious period in a fully susceptible population [1], can readily be written as *R*_0_ = *λ/µ* for this model. The mean duration of infection can be calculated as *δ* ∼ 1*/µ*. The likelihood of observing a certain transmission tree knowing the BDS model parameters has a closed-form solution. It is hence possible to estimate posterior distributions using a Bayesian framework [33] (see also Appendix 1).

In the following, we focus on epidemics caused by viruses because these are the most rapidly evolving microbes. The evolution of the virus genetic sequence within infected hosts is classically assumed to be decoupled from the transmission model and to have a fixed substitution rate. The likelihood of observing a certain multiple sequence alignment (MSA) at the sampled tips of a virus phylogenenetic tree (*i.e.*, genealogy) can be computed thanks to Felsenstein’s tree-pruning algorithm [12]. It is hence possible to infer a phylogenetic tree for a given MSA in a Bayesian [36, 4] or a maximum-likelihood framework [22, 26]. As viruses evolve rapidly, their sampling dates can be used to time-scale the phylogenetic tree [39, 32]. In the phylodynamics literature, the resulting time-scaled phylogenetic tree serves as a proxy for the transmission tree.

Following most phylodynamic studies, our datasets, namely MSAs, are simulated in two steps. First, the transmission trees are simulated under the BDS model using Gillespie’s algorithm [15]. To speed up the simulation, we reparameterize the BDS model by trans-forming the triplet (*λ, µ, s*) into 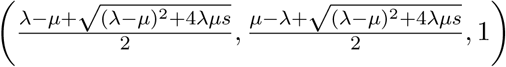, a method also used by Ref. [6]. This parametrization produces a sampled transmission tree, which is equivalent to the sampled transmission tree generated under the original model (the two differ in the unsampled parts but these are not used in our procedure). Second, multi-ple sequence alignments are simulated on these trees using AliSim [40], assuming a given substitution rate and a sequence evolution model.

### Simulated and real datasets

#### Simulated data for training and testing Teddy_0_

We define the prior distribution over the parameters used for training and test dataset gen-eration as follows: *R*_0_ ∼ U(1, 5) (where U denotes the uniform distribution), *δ* ∼ U(1, 10), nucleotide sequence evolution follows a Jukes-Cantor (JC) model [20], and the substitution rate follows a strict clock with the rate parameter is drawn from log_10_ U(−5, −3). The number of leaves is drawn uniformly between 20 and 100, and the sampling proportion is drawn from log_10_ U(−2, 0).

The generation of sequences along the transmission tree requires an initial sequence. It is randomly generated according to the equilibrium distribution of the evolution model (uniform over the four nucleotides in the case of JC model).

We generated 1 500 000 couples (parameters, dated MSA) for the training set and 1, 000 samples for the test set.

#### Real virus outbreak data

We used the data from a Hepatitis C Virus (HCV) outbreak [18], which consists in an MSA of 40 sequences of the viral gene coding for the nonstructural protein 5B (NS5B). These sequences were sampled from a cohort of people living with HIV in Bangkok, Thailand. We extracted sampling dates corresponding to these sequences from GenBank using their accession numbers. These ranged between March 25, 2015 and July 6, 2019. To make sure that the assumptions of the epidemiological and evolutionary models used in Teddy were applicable, we removed the gap positions from the alignment. Furthermore, to detect potentially problematic sequences, we performed a maximum likelihood phylogenetic tree reconstruction using RaxML-NG [22] and time-scaled it and detected temporal outliers with LSD2 [39]. This led to the identification and removal of one temporal outlier sequence (*i.e.*, a sequence whose substitution rate was more than 3 standard deviations higher than the mean one) and four sequences that belonged to a separate clade different from the main one. We chose to focus on one, very recent (starting in 2013), clade instead of keeping both clades, as the most recent common ancestor of the two clades dated from 1973, hence violating the BDS hypothesis of homogeneous sampling (the first sample dates from 2015). In the end, we retained 35 sequences for our analysis, annotated with sampling dates represented as real values (e.g., July 6, 2019 was represented as 2019.51).

#### Simulated data for training and testing Teddy_HCV_

For the real data, the prior distributions were adjusted based on prior knowledge about HCV epidemics. The prior for *R*_0_ was unchanged and we assumed that *δ* ∼ U(0.5, 10). Furthermore, we set the substitution rate to that estimated for another HCV outbreak, namely 1.3 · 10^−3^ substitutions/site/year [8]. We also assumed that sequence evolution followed the same model as in this previous work, namely a General Time Reversible (GTR) model [37] with gamma site heterogeneity model with 4 categories [46] and a proportion of invariant sites, with the following values: GTR{0.07349,0.817,0.0165,0.02485,1.0,0.0262} + G4{0.508}+I{0.499}.

The number of samples was drawn uniformly between 20 and 100, and the sampling proportion was drawn from log_10_ U(−2, 0).

The initial sequence for generating sequence evolution along the simulated trees was set to one of the earlier sequences in the real HCV dataset, namely 1a.TH.2015.HIVNAT 15.05.MN807647 (see the subsubsection on real outbreak data). We made this choice to ensure that the proportions of different nucleotides in the simulated MSAs resembled those of the real data.

As for the generic model, we generated 1 500 000 (parameters, dated MSA) couples for the training set and 1, 000 samples for the test set.

### Details on the Teddy architecture

We exploit the ideas of Geometric Deep Learning [5] that recommend encoding known symmetries of the estimated function in the neural network architecture to facilitate the learning procedure. More precisely, restricting oneself to sets of functions that verify as many known symmetries as possible makes the hypothesis class smaller, which reduces the estimation error of the procedure without increasing its approximation error. In our case, we know that the posterior of the epidemiological parameter is unaffected by two types of permutations over the input multiple sequence alignment (MSA). First, the order of the sequences in the MSA is arbitrary. Therefore, we want the output to be invariant under sequence permutation. Second, in both the JC and GTR models, as in most classical sequence evolution models, the sites are assumed to be independent from one another. Therefore, our estimated posterior should be identical if the sites are permuted. To achieve this double invariance, we use an axial transformer-like architecture, which combines the use of axial attention for equivariant transformations and a pooling technique based on added tokens for invariant transformation. It is schematically represented in Fig. 1 and detailed in the sections below.

#### Data representation and preprocessing

Our input is a fasta file containing an alignment of *n* genetic sequences with *l* sites. Each sequence identifier contains information about the sampling date. We use this data to fill an array where each row corresponds to a sequence and each column to a site. All constant sites are removed and the number of remaining variable sites is denoted by *l*. If there are too many variable sites in the MSA (*l̃ > l_max_*), we subsample them to keep the maximum number of allowed variable sites (*l_max_*) and discard the rest.

At the beginning of each row, we add the sampling date and a special token referred to as ‘CLS’ [9]. Finally, we add an initial row of ‘CLS’ tokens and, at the first position (above the dates), the sampling proportion. Our deep-learning model takes as input this array of shape (*n* + 1) × (*l̃* + 2), as well as the three quantities (*n, l,* *l̃*). For batching and compilation purposes, a padding procedure is used with another special token ‘PAD’ such that all arrays have the same shape.

#### Encoding

For clarity, we omit the batch dimension in the following. The input is decoupled into three components: the dates, the data (*i.e.*, the sequences and ‘CLS’ tokens), and the sampling proportion. We create a mask from the data to find the ‘PAD’ tokens. We use one-hot encoding to obtain a tensor of shape (*n* + 1) × (*l̃* + 1) × *d̃*. We set *d̃*= 11, corresponding to the 4 nucleotides, 1 gap, 2 special tokens (‘CLS’ and ‘PAD’), and the extra 4 pieces of information (sampling proportion *s*, number of sequences *n*, number of initial sites *l*, and number of non-constant sites *l̃*). The sampling proportion and the information about the shapes are added to the data through their corresponding entries.

The first learnable function is then applied to this embedding. It consists of a fully connected linear layer, and is followed by a ReLU activation function and a layer normal-ization. This layer allows us to increase the embedding’s dimension to *d* = 64. Therefore, our data gets a shape (*n* + 1) × (*l̃* + 1) × *d*.

We then add the sampling date information to each row. Given the invariance of the posterior under translation of all the dates (see Appendix 1 for more details), we subtract the smallest date value from all dates. Next, we map the dates to a vector of dimension *d* = 64 thanks to a function similar to the positional encoding [41], but used here in a continuous way. More precisely, for a date *x*, we define the vector TimeEncoding(*x*) on each dimension by

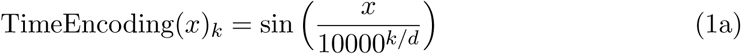

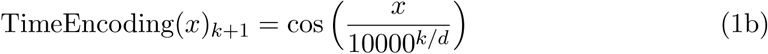

where *k* ∈ {0, 2, 4*, …, d*}. This encoding is added to the first column, which corresponds to the added ‘CLS’ column.

#### Axial transformers blocks

Our resulting (*n*+1)×(*l̃* +1)×*d* embedding is given as input to a series of axial transformer blocks.

In this work, we use 6 blocks, each of which is composed of the following layers:

1. **An attention layer on sequences**. In this step, we apply (*l̃* + 1) times the same attention heads on sets of (*n* + 1) embeddings, which allows the information to flow between all sequences at each site.
2. **An attention layer on sites**. In this step, we apply (*n*+1) times the same attention heads on sets of (*l̃* + 1) embeddings, which allows the information to flow between sites within each sequence.
3. **A feed forward layer**. This final step consists of a simple hidden layer followed by a normalization layer (except for the last axial layer) with a GELU activation function.

Following this step, we still have an embedding of shape (*n* + 1) × (*l̃* + 1) × *d*, but each entry in the array now contains information about the whole dated MSA.

All our self-attention operations are implemented through the FlexAttention frame-work [10]. This makes the computation time and memory usage of Teddy scale linearly in both the number of sequences and sites. We handle MSAs of variable size within each batch—from 20 to 100 sequences and up to 250 non-constant sites—by padding to the largest one and masking the padding elements in self-attention.

#### Projection and regression

From our last embedding, we extract the corner ‘CLS’ (*i.e.*, the intersection of the row ‘CLS’ and column ‘CLS‘). Because of the axial transformer layers, this vector of size *d* contains information about the whole dated MSA. Until this step, our transformation was equivariant to permutations of the sequences or sites, *i.e.*, permuting the sequence or sites in the input MSA led to the same permutation on the output of the transformation. The projection step defined by extracting the corner CLS makes the transformation invariant under the permutation of the sites and the sequences—the output of the projection is unaffected by any such permutation of the input MSA. We then use a feed-forward layer with a single hidden layer and a softplus activation function. The output has 6 components, and we cumulatively sum the first three and the last three. These resulting 6 components represent the estimate of the 0.025, 0.5 and 0.975 quantiles of the marginalized posterior distributions of *R*_0_ and *δ*.

The combination of the embedding, axial transformer, projection and regression layers described above defines a (learnable) function *f_ϕ_* that takes as input a dated MSA, here denoted *X*, and a sampling proportion *s*, and returns a vector

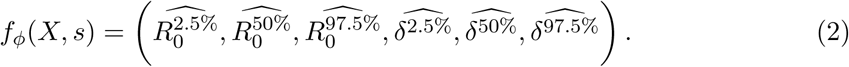

### Training procedure

#### Loss function

We train our networks to minimize a Pinball loss [35], which is known to be minimized by quantiles. We focus on the three quantiles of interest (0.025, 0.50, and 0.975) for each of the two parameters, namely *R*_0_ and *δ*. We then sum these six losses to obtain our total loss, which can be written as

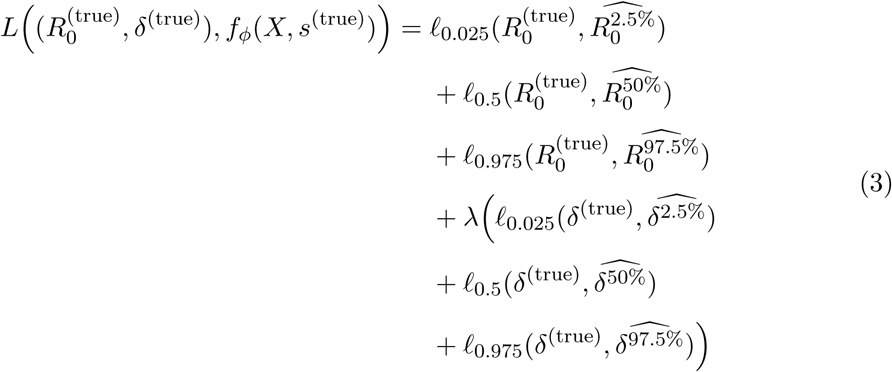

where *λ* ∈ *R*^+^ is a parameter set to give more importance in learning to either one of the parameters. We set *λ* = 5 in our experiments, as learning *δ* seemed to be more difficult than learning *R*_0_.

Before computing the loss, we normalize the outputs and the parameters of interest in order to have comparable loss values. To do so, we use an affine transformation such that outputs can also be compared. For the *R*_0_ parameters, we use the transformation 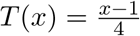 such that [1, 5] is sent to [0, 1]. For the *δ* parameter, we use 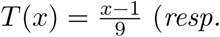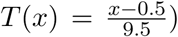 for Teddy_0_ (*resp.* Teddy_HCV_) such that [1, 10] (*resp.* [0.5, 10]) is sent to [0, 1].

Note that this normalization does not affect the gradient, as this one is piecewise constant and only depends on the sign of the difference between the output and the parameters. For that reason and readability, these transformations are not shown in the equation 3.

#### Optimizers, hyperparameters, and scheduler

To train Teddy_0_, we used AdamW [24] as optimizer with a weight decay of 0.01 and betas (*β*_1_*, β*_2_) = (0.9, 0.98). We used a cosine with linear warm-up learning rate scheduler, set such that the maximum learning rate was 8 × 10^−5^ reached after 10,000 steps. The maximum number of steps was set to 500,000 steps. We also limited the training time to 19 hours. The results shown originate from a model that ran 333,094 steps in 19 hours, corresponding to 58 epochs. We used a batch size of 64 for this experience, and a maximal number of variable sites *l_max_* to 250.

#### Finetuning

Note that we did not freeze any parameters of the model, as the early layers encode data, and invariance under permutation of nucleotides is lost between the two.

We finetuned from Teddy_0_ during 100, 000 steps during 469 minutes. We used the same optimizer, with a cosine with linear warm-up learning rate scheduler, set such that the maximum learning rate was 4 × 10^−4^ reached after 1, 000 steps. The batch size for this experience was set to 128, and *l_max_* to 180.

### Metrics

In our experiments, we analyze the Equal-Tailed credible Intervals (ETI), which are given by the quantiles at 2.5 % and 97.5 % of the posterior distribution.

Other key metrics are the MAE and the MRE, which were computed with respect to the inferred median of the posterior—*i.e.*, they measured how far over the 1, 000 alignments the inferred median was, on average, from the actual parameter used to sample the alignment.

### BEAST 2 parameterisation

We used the package bdsky [33] to model a homogeneous constant Birth-Death-Sampling model by assuming a Birth Death Skyline model with no changepoint.

For the Teddy_0_ simulated dataset, we assume the following priors:

- *R*_0_ ∼ U(1, 5), initial value: 3,
- removal rate 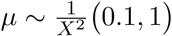, initial value: 0.18,
- strict clock substitution rate 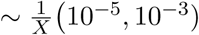, initial value: 10^−4^.

Here, 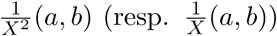 denotes the distribution with density 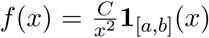 for all 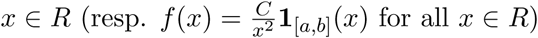 if 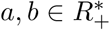 such that *a < b* and *C* ∈ *R_+_*^∗^ is a constant of normalization. This ensures the desired 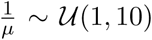 distribution. The implementation of such a prior is provided as an external package we coded and provide on the associated gitlab repository.

We set the sampling proportion to the value used for generating the corresponding dated MSA. For all samples, we set the evolution model to JC.

For the Teddy_HCV_ simulated dataset, we assume the following priors:

- *R*_0_ ∼ U(1, 5), initial value: 3,
- removal rate 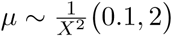, initial value: 0.19

We fixed the evolution model to GTR{0.07349,0.817,0.0165, 0.02485,1.0,0.0262}+G4{0.508}+I{0.499} and a constant clock rate to 1.3 · 10^−3^.

## Data, Materials, and Software Availability

The code associated with Teddy is available at https://gitlab.in2p3.fr/vincent. garot/teddy_official. All the materials used to generate the results shown in the arti-cle will be made available through the https://zenodo.org platform with a DOI before publication.

## Acknowledgments

This work was funded by the Agence Nationale de la Recherche (grant DEELOGENY ANR-23-CE45-0027 to LJ, SA, and AZ) and the DIM One HEALTH PhD fellowship (to VG). The authors were granted access to the high-performance computing (HPC)/AI resources of Institut du développement et des ressources en informatique scientifique (IDRIS) under the allocation AD011011137R1 made by the Grand Équipement National de Calcul Intensif (GENCI). They also acknowledge the ISO 9001 certified IRD i-Trop HPC (South Green Platform) at IRD Montpellier for providing HPC resources that have contributed to the research results reported within this article.

