## Supplementary Files for "Teddy: neural inference of epidemiological parameters from viral sequences"

### Supplementary Files for
### Teddy: neural inference of epidemiological parameters from viral sequences

Vincent Garot, Luc Blassel, Luca Nesterenko, Anna Zhukova, Samuel Alizon, Laurent Jacob

Corresponding Authors: Vincent Garot & Laurent Jacob
 &

### Supplementary Text S1: Relationship between epidemiological and evolutionary parameters

#### Relationship between time-scale and infectious duration

In this appendix, we show that a neural network trained over transmission trees for a given time scale of infection duration (*e.g.*, years) can be used to produce inference over epidemics occurring at different time scales (*e.g.*, months) by simply rescaling the observed sampling dates.

Let us consider a transmission tree $\mathscr{T}$ generated by a Birth-Death model with incomplete sampling (BDS) [4] with the rate parameters $\Theta = \{\lambda, \mu, s\}$ (birth (transmission) rate, death (becoming-non-infectious or removal) rate and sampling probability), which can also be represented as epidemiological parameters $\{R_0 = \frac{\lambda}{\psi}, \delta = \frac{1}{\psi}, s\}$, *i.e.*, (basic reproduction number, infectious duration, sampling probability). The tree loglikelihood formula under this model can be written as:

$$\log L(\mathscr{T}|\Theta) = \sum_{i \in \text{tips}} \log(\mu s \Delta t) \leftarrow \textit{sampling of tips} \quad (1)$$

$$+ \sum_{i \in \text{internal nodes}} \log\left(2\lambda \Delta t\right) \leftarrow \textit{transmission events} \quad (2)$$

$$+ \sum_{i \in \text{internal nodes}} \log\left(p^{(i_0)}(t_i) p^{(i_1)}(t_i)\right) \leftarrow \textit{node i's child branch evolution} \quad (3)$$

$$\text{where } p^{(i)}(t) = \left(\frac{E(t_i)+1}{E(t)+1}\right)^2 e^{c_1(t-t_i)},$$

$$c_1 = \sqrt{(\lambda-\mu)^2 + 4\lambda\mu s},\ c_2 = \frac{c_1+\lambda-\mu}{c_1-\lambda+\mu}, E(t) = c_2 e^{c_1(t-T)},$$

$t_i$ is the time at node $i$,

$T = \max_i t_i$ (time at the end of the sampling interval),

$i_0$ and $i_1$ are child nodes of $i$,

$\Delta t$ is an infinitely small time interval.

Note that the parameters $\lambda$ and $\mu$ have the dimension [1/time unit] (where the time unit is, for example, year), while the parameter $s$ does not depend on time (is dimensionless). The constant $c_1$ has the dimension [1/time unit], the constant $c_2$ is dimensionless, the formula $e^{c_1(t-T)}$ is dimensionless ([1/time unit] and [time unit] cancel out) and so are $E(t)$ and the branch evolution probabilities $p^{(i)}(t)$. The elements of the first two sums in the formula (3) are dimensionless as they include rates $[\frac{1}{\text{time unit}}]$ multiplied by time

[time unit] (dimensions cancel out), and the elements of the third sum are dimensionless probabilities. Hence, the tree likelihood does not depend on the chosen time units.

Let us consider a transmission tree with 3 sampled pathogen sequences: c, d, and e (as in the left panel of Fig. S6). We assume that this tree was generated by a birth-death-sampling (BDS) model with basic reproduction number $R_0$, infectious duration $\delta$ [years], and sampling probability $s$. This tree, which is scaled in years, has exactly the same likelihood as a tree with the same topology and the same relative branch lengths generated under parameters $R_0, \delta$ [months] and $s$, *i.e.*, on a month scale (as the likelihood formula (3) is dimensionless). Moreover, this tree also has the same likelihood if considered on the month scale (as in the right panel of Fig. S6(right)), with parameters $R_0, 12 \cdot \delta$[months] and $s$—as this just amounts to multiplying all branches and times by a factor 12 and dividing all the rates by a factor of 12 (which will cancel one another) in the likelihood formula (3).

Hence, any binary transmission tree $\mathcal{T}$ with $n$ tips, branch lengths $t_1$, $t_2$, ..., $t_{2n-1}$ [years], and infection duration value $\delta$ for the BDS model define an equivalence class of other (tree, duration) leading to the exact same probability distribution for all BDS parameters:

1. (equivalent trees on a different time-scale) all transmission trees $\mathcal{T}'$ with the same topology and branch lengths $t'_1 = t_1, t'_2 = t_2, \ldots, t'_n = t_n$ [months], $R'_0 = R_0$, $\delta' = \delta$ [months], and $s' = s$. The same holds for other time-scales (weeks, days, etc.)

2. (different time-scales for the same tree) all transmission trees $\mathcal{T}'$ with the same topology and branch lengths $t'_1 = 12t_1, t'_2 = 12t_2, \ldots, t'_n = 12t_n$ [months], $R'_0 = R_0$, $\delta' = 12\delta$ [months], and $s' = s$. The same holds for other time-scales (weeks, days, etc.)

Note that it is the branch lengths and not the tip sampling dates that are used in the likelihood formula (3). Hence, the sampling dates can be expressed relative to any event (*e.g.*, to the date of the tree root, like in Fig. S6, or to the beginning of the current era (AD), or to the sampling date of the oldest sampled tip). In our case, we can generate all the training data under $\delta$ at some arbitrary time scale, and use the training model for inference on any dated MSA by simply converting its sampling dates. We express sampling dates relative to the date of the first sample; and the time-scale such that $\delta \in ]1, 10]$.

In practice, the choice of the time-scale could also imply truncating sampling dates (*e.g.*, to months: Aug 2024 *vs.* days: 18 Aug 2024), and hence decreasing the resolution of the transmission tree. From this point of view, more precise dates should be preferred [1].

#### Relationship between pathogen genome substitution rate and alignment size

In our case, the input data is represented by the variable sites extracted from the aligned sampled sequences and their dates instead of time-trees. Variable sites correspond to positions where the character (nucleotide or amino acid) is different for at least two samples. In classical phylogenetics, the inference of a time-tree from a multiple sequence alignment and its sampling dates uses two additional models: the model of sequence evolution (here fixed to JC [3]) and the clock model. Both can be inferred from the input data. The strict molecular clock model assumes homogeneous substitution rates among the branches of the phylogeny. This model has a single parameter, the substitution rate ($r$), which is typically measured in substitutions per [alignment] site per unit of time (*e.g.*, years). Time-scaling converts a phylogenetic tree reconstructed from pathogen sequence data (with branches measured in substitutions per site) into a time-tree whose branches are measured in units of time, using the sampling dates of the tree tips [2]. The latter tree can then be used as a proxy of a transmission tree for epidemiological parameter inference in classical phylodynamics.

Under the strict clock model, the relationship between the length $t_i$ of a branch $i$ in the time-tree and the length $b_i$ of the corresponding branch in the corresponding phylogenetic tree can be represented as: $b_i = r \cdot t_i$ (in practice, the right-hand side also includes a summand $\epsilon_i$ representing a noise term accounting for sampling and inference errors, and discrete nature of the substitution-per-site measure *vs.* continuous nature of time).

This (ideal) formula implies that for the same transmission tree such as the one in Fig. S6 (on the left on the year scale and on the right on the month scale), we expect a pathogen $p_1$ with a substitution rate $r_1$ to have branches in its phylogenetic tree with $n$ times less substitutions than the corresponding branches in the phylogenetic tree of a pathogen $p_2$ whose substitution rate $r_2$ is $n$ times higher: $r_2 = n \cdot r_1$ (on the same time scale). Provided they have the same alignment lengths $A_1 = A_2$, and a "reasonable" (as explained in the next subsection) sampling interval, this implies that the $p_1$'s alignment is expected to have $n$ times fewer variable sites. For instance, in Fig. S6, $r_2 = 2 \cdot r_1$ and the alignment on the left has half as many variable positions as the alignment on the right (8 *vs.* 16). However, if $p_1$'s alignment length is $n$ times longer ($A_1 = n \cdot A_2$, *e.g.*, if we only sequenced the first half of the genome for the pathogen on the right in Fig. S6: positions 1-9), this would translate to the same number of variable sites (8). Hence, the increase in the substitution rate and/or in the sequenced genome part length adds "resolution" to the phylogeny branch lengths (until saturation is reached, see the next subsection). It does not, however, change the underlying transmission tree.

Therefore, as long as the pairwise common-to-different variable site proportions $f_{ij}$ in sampled sequences stay (almost) the same (*e.g.*, d *vs.* e have $f_{de} = 3/5$ (positions 2, 3, and 11 *vs.* positions 4, 6, 12, 15 and 17) in the alignment in the left panel of Fig. S6 and $f_{de} = 6/10 = 3/5$ in the right panel; etc.), and the sampling dates stay compatible (as discussed above), the two datasets belong to the same equivalence class. Hence, to reduce the training data set, only one of the two parameters (sequence length, substitution rate) needs to be varied.

#### Settings where estimation works and where it might not

For a transmission order to be identifiable from the genomic data, at least one substitution is needed between two consecutive transmissions (*i.e.*, along an internal branch of a transmission tree). The average length of an internal branch in a complete transmission tree (including sampled and unsampled parts) can be approximated with $1/\lambda$, where $\lambda$ is the transmission rate. Having at least one (expected) substitution along this branch would correspond to a substitution rate $r \geq \lambda/l$, where $l$ is the alignment length.

On the other hand, having too many substitutions may lead to saturation and erase the evolutionary history. Saturation would correspond to $\geq 1$ substitutions per site between two observed events (*e.g.*, transmissions, *i.e.*, along an internal branch in an observed transmission tree). For simplicity, we would instead consider a more conservative scenario of a pathogen genome reaching $\geq 1$ substitutions per site during the sampling period. The sampling period corresponds to the tree height $H$. Depending on the tree shape, the path between the root and the most recently sampled tip can include between $O(logN)$ (balanced tree) and $O(N)$ (ladder-like tree) branches, where $N$ is the number of tips in the complete transmission tree. Hence, in the most conservative case (ladder-like tree), we can approximate the tree height $H \approx N/\lambda$. Hence, to avoid saturation, the substitution rate $r$ should be $\leq \lambda/N$. Note that if the alignment contains $n$ samples at a sampling proportion of $s$ and the reproduction number is $R_0$, then $N = nR_0/s$.

Therefore, we can define a "reasonable" setting for estimation of BD epidemiological parameters from pathogen sequence data, as the one where the substitution rate $r \in [\frac{\lambda}{l}, \frac{\lambda s}{nR_0}]$, where $l$ is the alignment length in terms of the number of sites and $n$ in terms of the number of sequences. $r < \frac{\lambda}{l}$ corresponds to limited phylogenetic signal, and $r > \frac{\lambda s}{nR_0}$ to potential saturation.

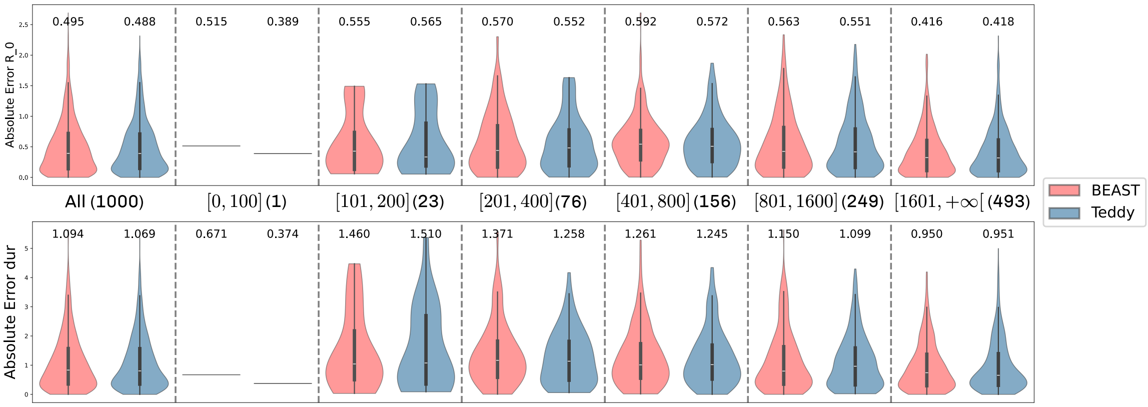


Figure S1: **Distribution of the Mean Absolute Error (MAE) as a function of the ESS of the BEAST 2 MCMC.** The ESS was taken to be the lowest one between that of $R_0$ and $\delta$. The MAE was computed on $1,000$ samples under the BDS-JC model described in the Methods. The numbers in parentheses correspond to the number of samples in each bin. The number above each violin plot indicates the mean of the distribution.

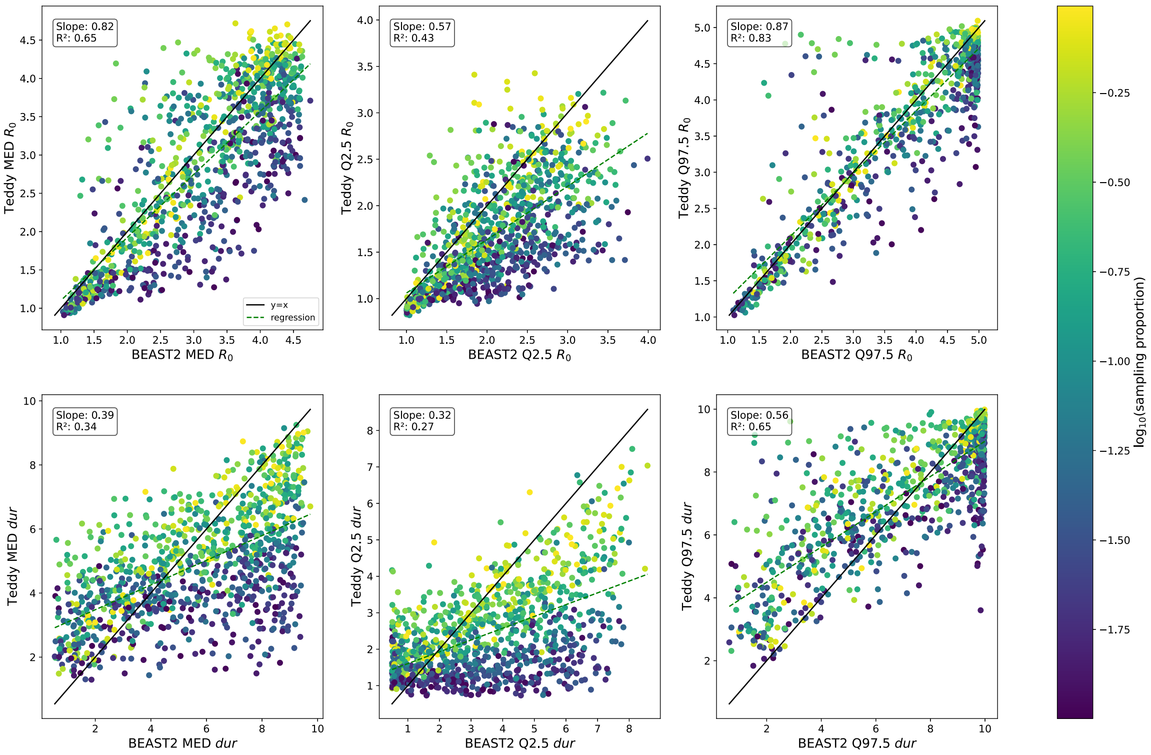


Figure S2: **Estimations of Teddy$_0$ versus BEAST 2 for the quantiles using the HCV simulated data.** Each point corresponds to the estimate on one of the 1,000 test MSAs generated under the HCV-like BDS-GTR model described in the Methods. See Figure 2 in the main text for details.

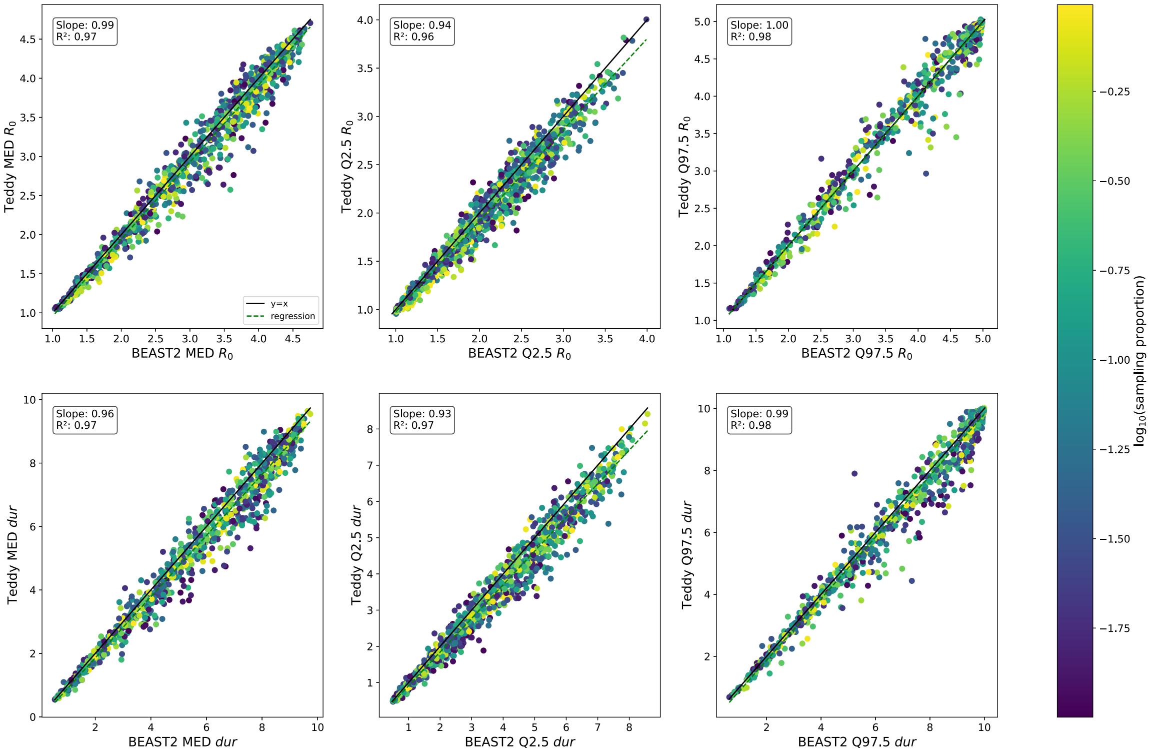


Figure S3: **Estimations of $\text{Teddy}_{\text{HCV}}$ versus BEAST 2 for the quantiles using the HCV simulated data.** See Figure S2 for details.

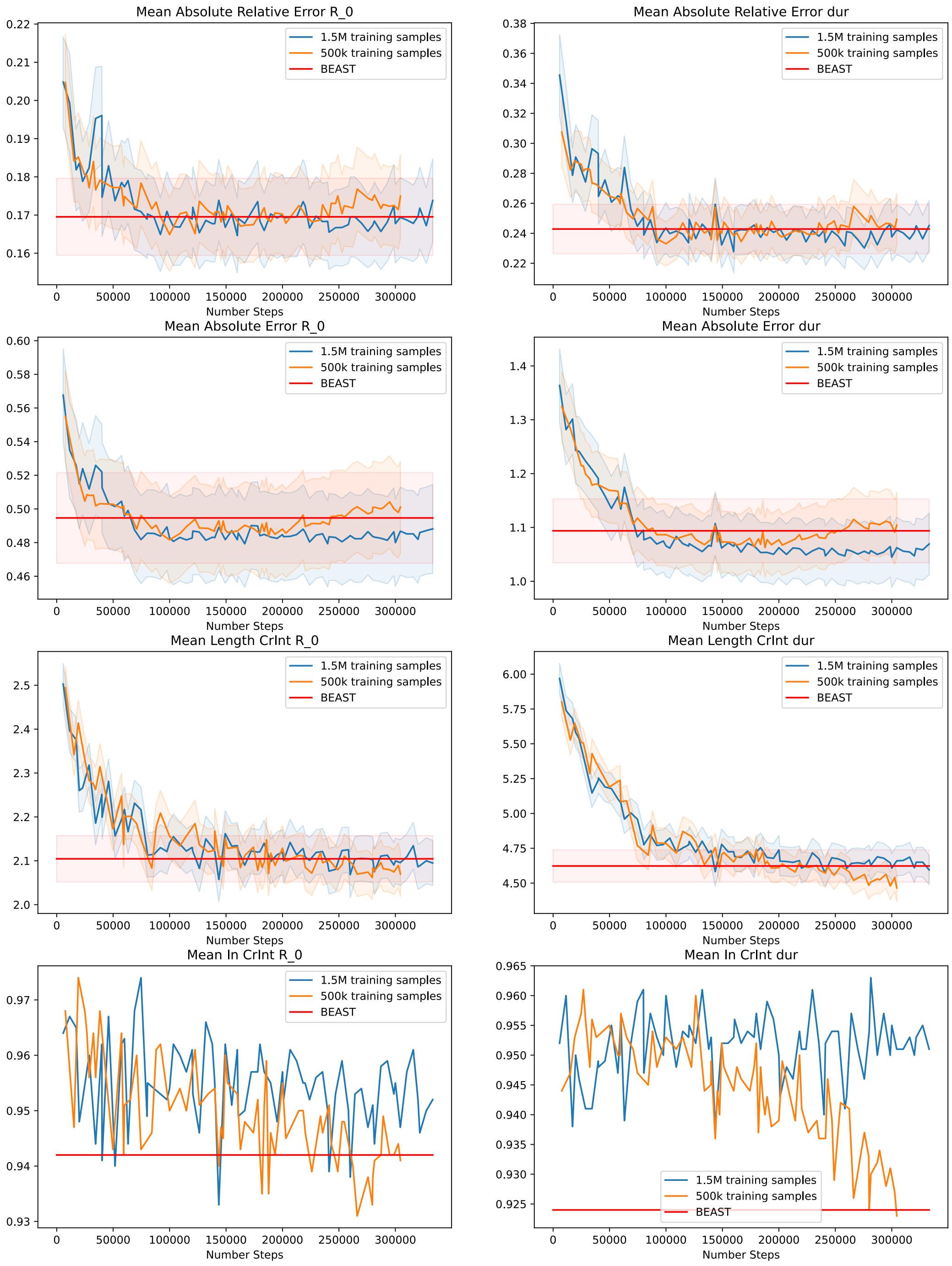


Figure S4: Validation metrics along training steps for identical architectures and training hyperparameters using 500, 000 (orange) or 1, 500, 000 (blue) training examples. The shaded intervals correspond to 1.96 standard errors.

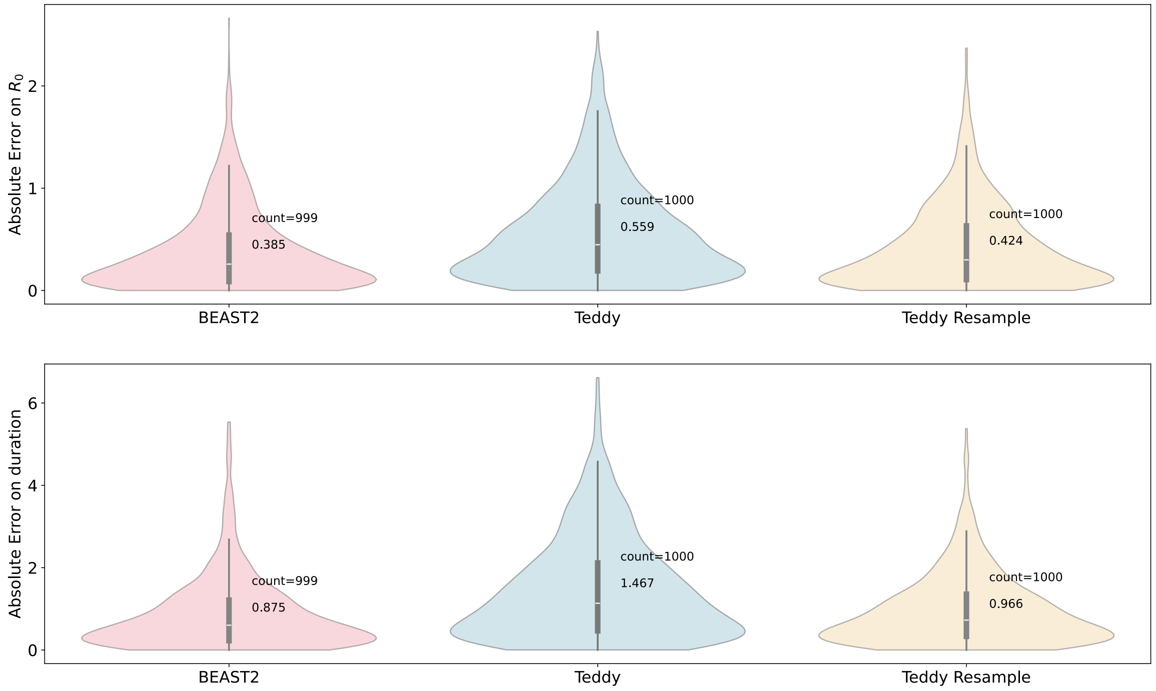


Figure S5: **Analyzing large sequence alignments with BEAST 2, $\mathbf{Teddy_0}$, and an ensembling version of $\mathbf{Teddy_0}$.** Each of the three methods is applied to 1,000 MSAs of 200 sequences generated under the model described in the Methods. The ensembling version is denoted 'Teddy resample' and draws 10 subsets of 100 sequences from the 200, infers parameters with $\text{Teddy}_0$ (denoted 'Teddy') on each subset, and returns the median of the estimates. The top and bottom panels show the distribution of the 1000 median Mean Absolute Errors (MAE) for estimates of $R_0$ and $\delta$, respectively. One run of BEAST2 has been discarded due to non-convergence.

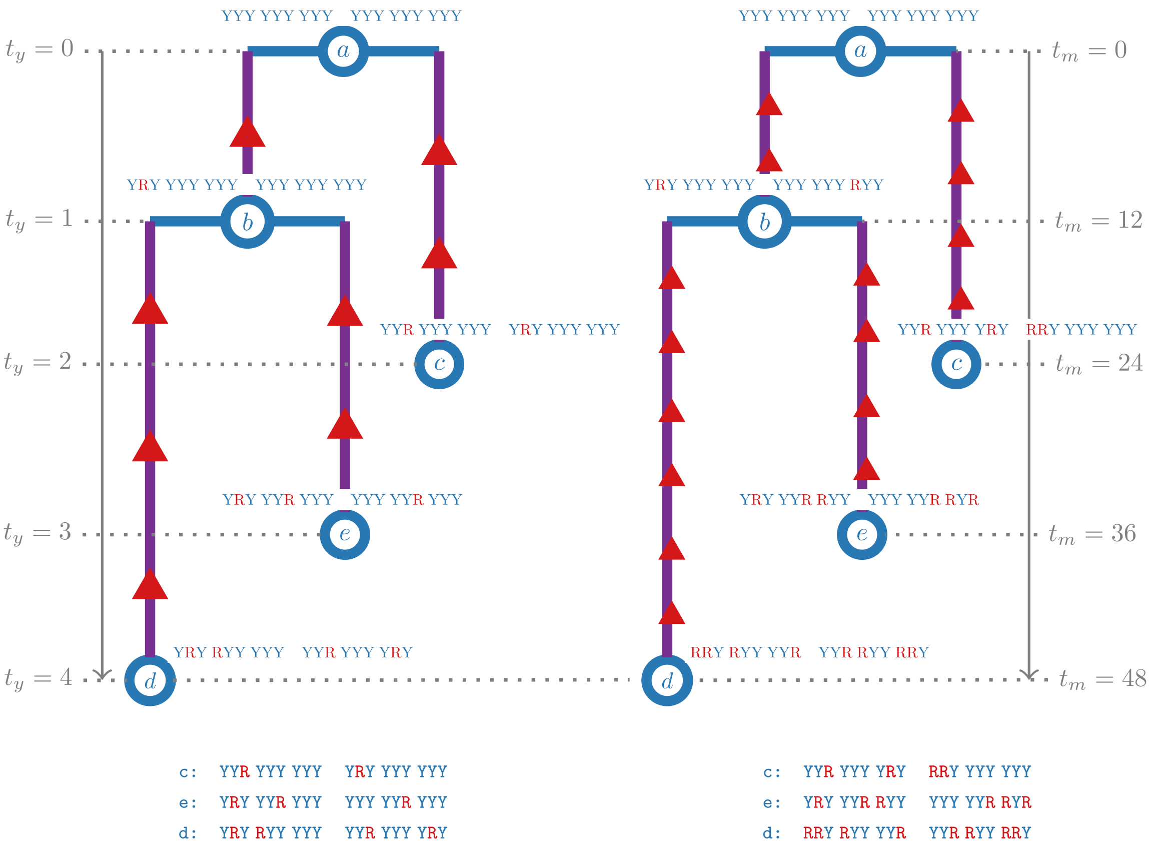


Figure S6: **Relationship between transmission trees, substitution rate and sequences** Both panels represent a transmission tree for 3 sampled sequences: c, e, and d. On the year time-scale (left) c is sampled at year 2, e is sampled at year 3, and d is sampled at year 4, while on the month time-scale (right) they are sampled correspondingly at month 24, 36, and 48. Their genomic sequences, as well as the ancestral sequences, are shown on top of the corresponding tree nodes. The corresponding multiple sequence alignments (of length 18 [sites]) are shown below each tree. The tree on the left corresponds to the substitutuon rate $r_1 = \frac{1}{18}$ [substitutions per site per year] = $\frac{1}{18 \cdot 12}$ [substitutions per site per month], while the tree on the right – to the twice higher substitutuon rate $r_2 = 2r_1$. The substitutions that occurred over time are shown as right triangles on the corresponding branches. For simplicity, we consider only two types of genomic characters: purine (R) vs. pyrimidine (Y), instead of 4 DNA characters or 21 amino acid characters. The alignment on the left contains 8 variable sites (2nd, 3rd, 4th, 6th, 11th, 12th, 16th, and 18th), while the one on the right contains twice as many (16: all but the 5th and the 11th) variable sites. Note that if for the second case, only the first half of the genome was sequenced instead (positions 1-9), the resulting alignment would have had 8 variable sites, as in the first case.

Table 1: **Average performance of BEAST 2, $\text{Teddy}_0$ and $\text{Teddy}_{\text{HCV}}$ on HCV-like simulated datasets.** Mean Absolute Error (MAE) and Mean Relative Error (MRE) are computed over 1 000 test MSAs. The coverage corresponds to the percentage of true parameters in the predicted 95% Equal-Tailed credible Interval (ETI). We also compute the Mean Width of the estimated ETI over the test set. For the MAE, MRE and the Mean Width ETI, we compute the standard error (ste) of the estimator of the mean. The values corresponding to the best performance are shown in bold.

| | $R_0$ | | | $\delta$ | | |
|---|---|---|---|---|---|---|
| | BEAST 2 | $\text{Teddy}_0$ | $\text{Teddy}_{\text{HCV}}$ | BEAST 2 | $\text{Teddy}_0$ | $\text{Teddy}_{\text{HCV}}$ |
| MAE (ste) | 0.464 (0.013) | 0.698 (0.018) | **0.458** (0.013) | 0.865 (0.024) | 1.84 (0.046) | **0.836** (0.024) |
| MRE (ste) | 0.166 (0.005) | 0.248 (0.008) | **0.159** (0.005) | 0.215 (0.008) | 0.588 (0.028) | **0.195** (0.007) |
| % in ETI | 92.0 | 81.8 | **95.2** | 92.2 | 73.5 | **94.1** |
| Mean Width ETI (ste) | **1.91** (0.03) | 2.17 (0.02) | 1.99 (0.03) | **3.61** (0.05) | 4.98 (0.05) | 3.72 (0.05) |
